# The ecology of the Chloroflexi in full-scale activated sludge wastewater treatment plants

**DOI:** 10.1101/335752

**Authors:** Marta Nierychlo, Aleksandra Miłobędzka, Francesca Petriglieri, Bianca McIlroy, Per Halkjær Nielsen, Simon Jon McIlroy

**Affiliations:** Center for Microbial Communities, Department of Chemistry and Bioscience, Aalborg University, Aalborg, Denmark; Microbial Ecology and Environmental Biotechnology Department, Institute of Botany, Faculty of Biology, University of Warsaw; Biological and Chemical Research Centre, Żwirki i Wigury 101, Warsaw 02-089, Poland; Department of Biology, Faculty of Building Services, Hydro and Environmental Engineering, Warsaw University of Technology, 00-653 Warsaw, Poland

**Keywords:** Activated sludge, Filamentous bulking, Chloroflexi, *‘Candidatus* Amarolinea’, FISH, Microautoradiography

## Abstract

Filamentous bacteria belonging to the phylum Chloroflexi have received considerable attention in wastewater treatment systems for their suggested role in operational problem of impaired sludge settleability known as bulking. Their consistently high abundance in full-scale systems, even in the absence of bulking, indicates that they make a substantial contribution to the nutrient transformations during wastewater treatment. In this study, extensive 16S rRNA amplicon surveys of full-scale Danish WWTPs were screened to identify the most numerically important Chloroflexi genera. Fluorescence *in situ* hybridization probes were designed for their *in situ* characterization. All abundant phylotypes of the phylum were identified as facultative anaerobic chemoorganotrophs involved in fermentation of sugars. These groups were all filamentous but differed in their morphology and spatial arrangement. *‘Candidatus* Villigracilis’ was predominantly located within the activated sludge flocs, where they possibly have structural importance, and their abundance was relatively stable. Conversely, the abundance of *‘Candidatus* Amarolinea’ was highly dynamic, relative to other genera, sometimes reaching abundances in excess of 30% of the biovolume, suggesting their likely role in bulking episodes. This study gives an important insight into the role of Chloroflexi in WWTPs, thus contributing to the broader goal of understanding the ecology of these biotechnologically important systems.

## Introduction

Members of the phylum Chloroflexi constitute a substantial proportion of the activated sludge community in full-scale activated sludge wastewater treatment plants (WWTPs), where they reportedly constitute up to 30% of the biovolume and often make up the majority of filamentous bacteria present (Beer *et al*. 2006; Morgan-Sagastume, Nielsen and Nielsen 2008; Mielczarek *et al*. 2012). Filamentous bacteria are generally believed to have structural importance for activated sludge flocs with good settling properties. However, overgrowth of certain filamentous species is associated with open and diffuse flocs as well as interfloc bridging, leading to a sometimes severe operational problem known as bulking (Wanner, Kragelund and Nielsen 2010). Some Chloroflexi species have also been proposed to be involved in the stabilization of problematic foams at WWTPs (Kragelund *et al*. 2011) and membrane fouling in membrane bioreactors (MBRs) (Ziegler *et al*. 2016). In addition, the high abundance of the Chloroflexi in wastewater treatment systems, often in the absence of bulking problems, indicates that they make a proportional contribution to the observed nutrient transformations of these systems. As such, the study of this phylum has implications for plant operation and our general understanding of the ecology of wastewater treatment.

Relatively little is known about the ecology of the Chloroflexi in nutrient removal WWTPs. *In situ* studies reveal a preference for the uptake of sugars, and a high level of surface associated hydrolytic enzymes indicates their involvement in the breakdown of complex organics (Kragelund *et al*. 2007, 2011; Xia, Kong and Nielsen 2007). Due to poor phylogenetic annotation of the routinely applied taxonomic databases, studies of activated sludge community dynamics often consider members of the Chloroflexi phylum as a whole - an approach that ignores the likely phenotypic diversity among its members (McIlroy *et al*. 2015). An understanding of the ecology of the phylum and how it relates to system function requires the identification and *in situ* characterization of the abundant genus-level phylotypes present.

Historically, the identification of bulking filaments has relied on classification keys based on morphological characteristics, with most morphotypes identified by an Eikelboom number (Eikelboom 2000; Jenkins, Richard and Daigger 2004). More recent phylogenetic identification of these filaments has relied on 16S rRNA gene-based clone library analyses coupled with fluorescent *in situ* hybridization (FISH) of plants with severe filamentous bulking. Most of the antecedent morphotypes are possessed by members of the Chloroflexi and include: 1851 (Beer *et al*. 2002); 0092 (Speirs *et al*. 2009); 0803 (Kragelund *et al*. 2011; Speirs, Tucci and Seviour 2015); 0914 (Speirs *et al*. 2011); 0041/0675 (Speirs *et al*. 2017), and several others (Kragelund *et al*. 2009) - noting that organisms possessing the same filamentous morphotype are often unrelated (Seviour *et al*. 1997; Speirs, Tucci and Seviour 2015), and closely related organisms can also possess different morphotypes (Speirs *et al*. 2017). FISH probes available for known phylotypes reportedly cover many of the Chloroflexi present, but there is still a substantial portion of the phylum without genus-level probes (up to 90% by FISH in some plants) (Kragelund *et al*. 2011). The recent extensive MiDAS 16S rRNA gene amplicon sequencing-based survey, covering >50 Danish full-scale WWTPs over a 10 year period, has given a comprehensive overview of the abundant core members of the full-scale activated sludge treatment plants (McIlroy *et al*. 2015). As such, we are now able to systematically target the numerically important Chloroflexi phylotypes. The *in situ* physiology of some of these groups, such as ‘*Candidatus* Promineofilum’ (the B45 group) (McIlroy *et al*. 2016) and P2CN44 (Kragelund *et al*. 2011), has been determined, but several abundant phylotypes are novel and known only by their 16S rRNA gene sequence (McIlroy *et al*. 2015).

The aim of this study is to determine the *in situ* physiology of selected abundant novel Chloroflexi activated sludge phylotypes. The extensive MiDAS survey of full-scale Danish nutrient removal activated sludge plants was used to identify the most abundant genus-level taxa belonging to the phylum Chloroflexi. FISH probes were designed for these phylotypes and applied in combination with microautoradiography (MAR) and histochemical staining for their *in situ* characterization.

## Methods

### Biomass sampling and fixation for FISH

Biomass samples from the aerobic stage of selected full-scale activated sludge WWTPs with nutrient removal were fixed with 4% paraformaldehyde (PFA) for 3 h at 4°C. After fixation, samples were washed 3 times in sterile filtered tap water, re-suspended in 50% ethanol in 1 x PBS solution [v/v], and stored at −20 °C. For basic operational information for WWTPs sampled in this study, see Mielczarek *et al*., (2013).

### Phylogenetic analysis and FISH probe design

Phylogenetic analysis and FISH probe design were performed with the ARB software package (Ludwig *et al*. 2004) with the MiDAS database (Release 2.1), which is a version of the SILVA database (Release 123 NR99) (Quast *et al*. 2013) curated for activated sludge and anaerobic digester sequences (McIlroy *et al*. 2015, 2017b). Potential probes were assessed *in silico* with the mathFISH software (Yilmaz, Parnerkar and Noguera 2011). The Ribosomal Database Project (RDP) PROBE MATCH function (Cole *et al*. 2014) was used to identify non-target sequences with indels (McIlroy *et al*. 2011). Probe validation and optimization were based on generated formamide dissociation curves (Daims, Stoecker and Wagner 2005), where average relative fluorescent intensities of at least 50 cells calculated with ImageJ software (National Institutes of Health, Maryland, USA) were measured for varied hybridization buffer formamide concentration in increments of 5% (v/v) over a range of 0-65% (v/v) (data not shown). Details for probes designed in this study have been deposited in the ProbeBase database (Greuter *et al*. 2016).

### FISH

FISH was performed as detailed by Daims *et al*., (2005) using the probes designed in this study as well as CFX197 and CFX223, targeting *‘Ca*. Promineofilum’ (Speirs *et al*. 2009); CFX1223 (Björnsson *et al*. 2002), and GNSB941 (Gich, Garcia-Gil and Overmann 2001), applied as a mix to target the phylum Chloroflexi; EUB-338-I, EUB338-II, and EUB338-III (Amann *et al*. 1990; Daims *et al*. 1999), applied as a mix (EUBmix) to cover all bacteria; NON-EUB as a negative control for hybridization (Wallner, Amann and Beisker 1993). The hybridization conditions applied for each probe are given in **Table 1** or as recommended in their original publications. Quantitative FISH (qFISH) biovolume fractions of individual Chloroflexi genera were calculated as a percentage area of the total biovolume, hybridizing the EUBmix probes, that also hybridizes with the specific probe. qFISH analyses were based on 25 fields of view taken at 630 x magnification using the Daime image analysis software (Daims, Lücker and Wagner 2006). Microscopic analysis was performed with an Axioskop epifluorescence microscope (Carl Zeiss, Oberkochen, Germany), an LSM510 Meta laser scanning confocal microscope (Carl Zeiss), and a white light laser confocal microscope (Leica TCS SP8 X).

**Table 1.**
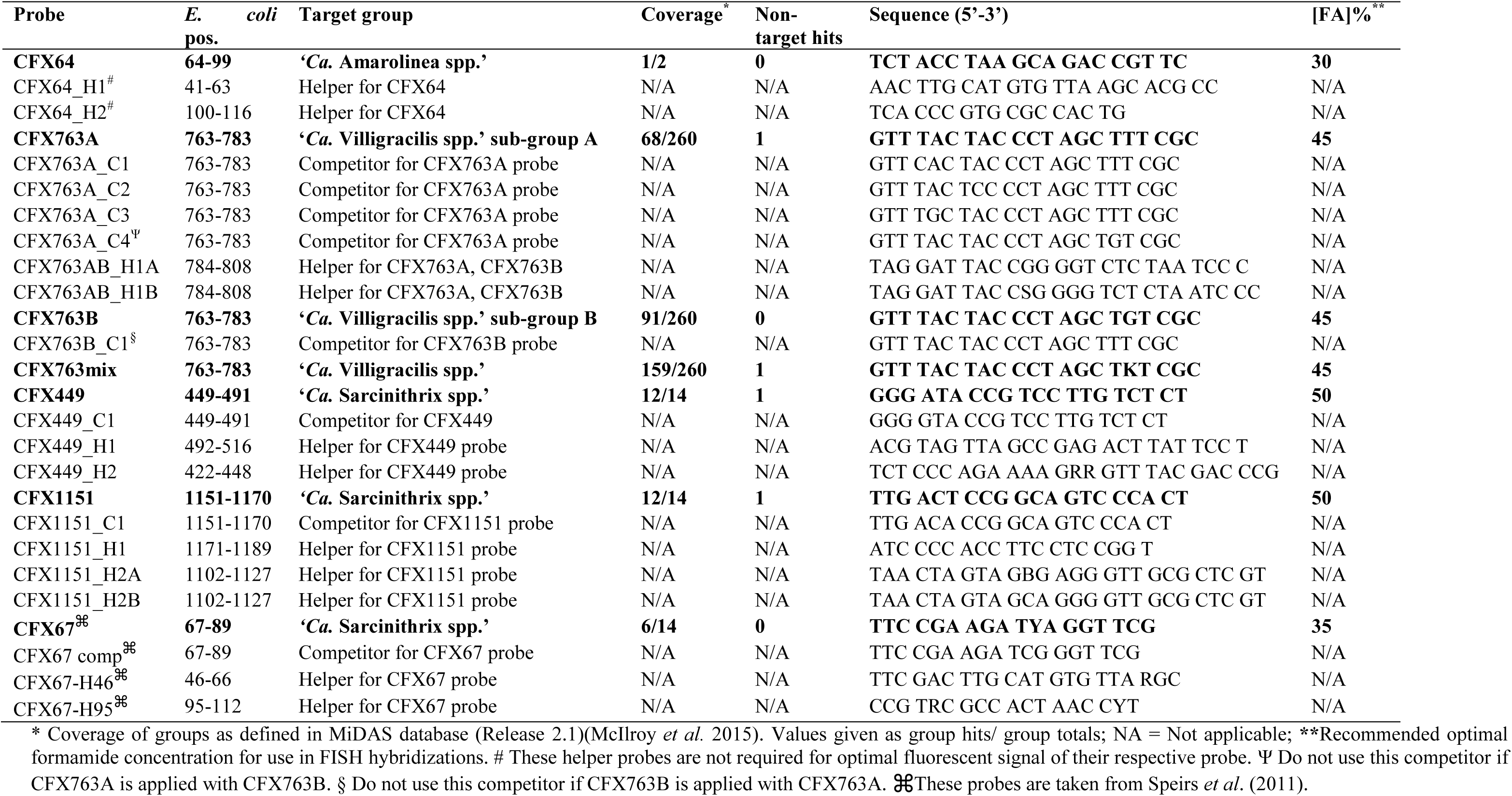
Probes designed and optimized in this study.

### Morphological classification

Microscopic observations of wet mount preparations and Gram and Neisser staining were performed according to the methods of Eikelboom (2000). Morphotype classification was carried out conforming to the described classification keys, based on morphological features of the filaments: shape, length, and diameter of filamentous bacteria, motility, presence of branching, attached growth of other bacteria to the filaments, (visible or not visible) septa between adjoining cells, shape of cells, presence of a sheath and sulphur granules (Eikelboom 2000).

### Histochemical staining

Following FISH, polyphosphate inclusions were stained with 28 μM 4’,6-diamidino-2-phenylindole (DAPI) for 1 h at 4°C in the dark. After staining at such relatively high DAPI concentrations, the polyphosphate granules appear bright yellow with fluorescence microscopy (Streichan, Golecki and Schon 1990). Polyhydroxyalkanoates (PHA) were stained with 1% [w/v] Nile Blue A for 10 mins at 55°C essentially as described previously (Ostle and Holt 1982). FISH images were acquired prior to staining and the same fields of view relocated.

### Microsphere adhesion to cells (MAC)

MAC was performed on a sample of fresh sludge to identify the hydrophobicity of target cells, applying the method of Kragelund and colleagues (2005) using a sonicated solution of 0.2 μm FluoSpheres fluorescent sulphate-modified microspheres with excitation and emission properties (505/515 nm) (Life Technologies Corporation, Eugene OR, USA).

### Microautoradiography

Biomass was sampled from the aerobic stage of full-scale activated sludge WWTPs in Aalborg West, Bjergmarken, Ringkøbing, and Odense North-West, Denmark. All plants are designed for N removal and enhanced biological phosphorus removal (EBPR) and have stable performance. For further details on the plants assessed in this study, see Mielczarek *et al*., (2013). Biomass samples were stored at 4°C and all incubations performed within 24 h from sampling. The MAR incubation protocol was based on the method of Nierychlo et al., (2015). Activated sludge was aerated for 40 min at room temperature prior to MAR incubation to reduce the residual substrates, oxygen, and NO_x_ present. Sludge was then diluted with filtered sludge water from the same plant to yield a biomass concentration of 1 mgSSmL^−1^ and transferred to 11 ml serum bottles. Radiolabeled substrates were added to yield a total radioactivity of 10 μCi mg^−1^ SS. The following substrates were used: [^3^H]acetate, [^3^H]galactose (Amersham Biosciences, UK), [^3^H]glucose, [^3^H]mannose, [^14^C]pyruvate (Perkin-Elmer, Waltham MA, USA), [^3^H]amino acid mix, [^14^C]butyric acid, [^3^H]fructose, [^3^H]glycerol, [^3^H]ethanol, [^3^H]lactate, [^3^H]NAG, [^14^C]propionate (American Radiolabeled Chemicals Inc., Saint Louis MO, USA). The corresponding cold substrate was added to yield a total concentration of 2 mM organic substrate. To achieve anaerobic conditions, prior to substrate addition, oxygen was removed by repeated evacuation of the headspace and subsequent injection of O_2_-free N_2_. Anaerobic incubations with selected substrates were supplemented with 0.5 mM nitrite or 2 mM nitrate to assess their use as electron acceptors. The supernatant concentrations were monitored using Quantofix Nitrate/Nitrite strips (Macherey-Nagel, Düren, Germany) and readjusted to their initial concentrations anaerobically to prevent exhaustion. Samples were incubated for 3 h at room temperature (approx. 21°C) on a rotary shaker at 250 rpm. Incubations with [^14^C]carbonate (American Radiolabeled Chemicals Inc., Saint Louis MO, USA) contained 20 μCi mg^−1^ SS of radiolabeled substrate. 1mM NH_4_Cl was added to half of the incubations to investigate ammonia and nitrite (produced from the oxidation of added ammonia) oxidation activity. These samples were incubated aerobically (same conditions as above) for 5h, as suggested by Daims et al., (2001). A pasteurized biomass (heated to 70°C for 10 min) incubation was prepared as a negative control to assess possible silver grain formation due to chemography. Incubations were terminated by the addition of cold PFA to a final concentration of 4% [w/v]. Samples were fixed for 3 h at 4°C and subsequently washed 3 times with sterile filtered tap water. Aliquots of 30 μl of the biomass were gently homogenized between glass coverslips. Following FISH (see earlier), coverslips were coated with Ilford K5D emulsion (Polysciences, Inc., Warrington, PA, USA), exposed in the dark for periods of 10 days, and developed with Kodak D-19 developer.

## Results

### Distribution of Chloroflexi in Danish full-scale WWTPs

Phylogenetic tree based on 16S rRNA gene sequences shows all the abundant Chloroflexi groups found in Danish WWTPs with nutrient removal and their phylogenetic relationship (**Figure 1**). The Chloroflexi phylum is among the most abundant phyla in full-scale systems in Denmark, constituting on average 10.6% of the total reads across all plants (**Figure 2A**). The Chloroflexi classes Anaerolineae, Caldilineae, Ardenticatenia, and SJA-15 made up the majority of members of the phylum present (**Figure 1B**). All the abundant genus-level phylotypes (**Figure 1C**) are novel, having no available cultured representatives, and were initially given provisional alphanumeric names in the MiDAS database (McIlroy *et al*. 2015) (not shown here). Based on their characterization, as presented in this study and previous publications, we propose new, previously unpublished, candidate names (Murray and Stackebrandt 1995). These names have been incorporated into the MiDAS taxonomy version 2.1 (McIlroy *et al*. 2015) and are used throughout this report. These include: ‘*Candidatus* Sarcinithrix’ (Sar.ci’ni.thrix. L. fem. n. *sarcina* a package, bundle; Gr. fem. n. *thrix* hair; N.L. fem. n. *Sarcinithrix* a hair bundle; formerly *Candidatus* Sarcinathrix (release 2.1)), ‘*Candidatus* Villigracilis’ (Vil.li.gra’ci.lis. L. masc. n. *villus* a tuft of hair; L. adj. *gracilis* slim, slender; N.L. fem. n. *Villigracilis* a slender tuft of hair; formerly MiDAS taxon SBR1029 (release 1.21) and *Candidatus* Villogracilis (release 2.1)), ‘*Candidatus* Defluviifilum’ (<u>De.flu.vi.i.fi’</u>lum. L. neut. n. *defluvium* sewage; L. neut. n. *filum* a thread; N.L. neut. n. *Defluviifilum* a thread from sewage; formerly MiDAS taxon P2CN44 (release 1.21)), and ‘*Candidatus* Amarolinea’ (A.ma.ro.li’ne.a. Gr. fem. n. *amara* conduit, channel, sewer; L. fem. n. *<u>linea</u>* a thread, a line; N.L. fem. n. *Amarolinea* a thread from a sewer; formerly MiDAS taxon C10_SB1A (release 1.21) and *Candidatus* Amarilinum (release 2.1)). ‘Kouleothrix spp.’, possessing the 1851 bulking filament morphotype, was present in low abundance with a median and mean of 0.04 and 0.4%, respectively. *‘Ca*. Defluviifilum’, *‘Ca*. Promineofilum’, *‘Ca*. Villigracilis’, and *‘Ca*. Sarcinithrix’ represent the four most abundant genera by median read abundance (**Figure 1C**), collectively constituting on average 6.2% of the total reads across all Danish plants assessed in this study. These phylotypes were relatively stably present across the different WWTPs (**Figure S1**) and therefore represent core members of the microbial community of these systems. As little is known regarding the physiology of the latter two genera, they were selected for a detailed characterization in this study. Relative to these phylotypes, the *‘Ca*. Amarolinea’ showed a much more dynamic distribution and periodically reached abundances in excess of 30% of the amplicon reads, which would indicate a likely role in bulking episodes in Denmark. As such, this genus, known only by its 16S rRNA gene sequences, was also selected for characterization in this study.

**Figure 1.**
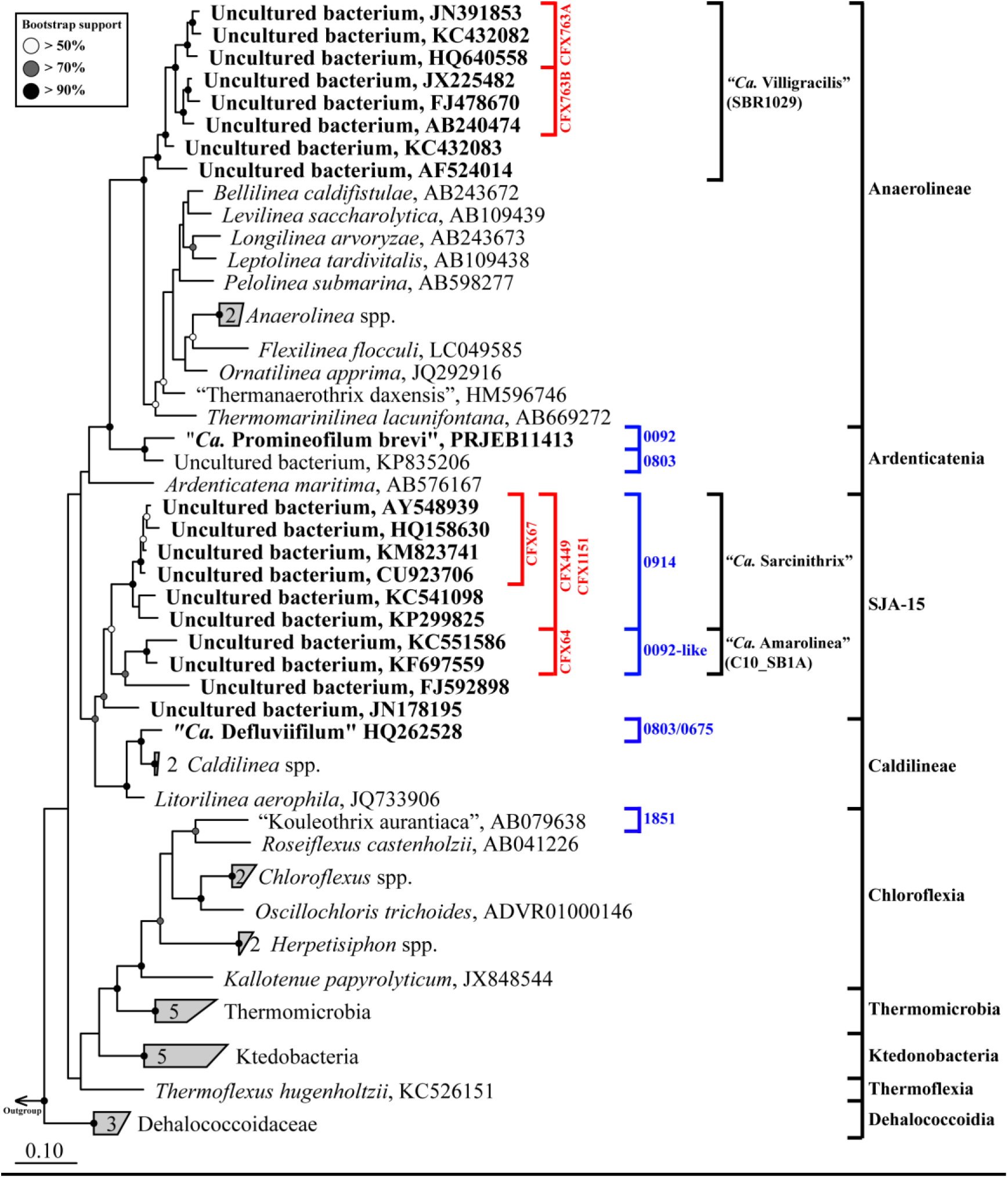
Maximum-likelihood (PhyML) 16S rRNA gene phylogenetic tree of abundant activated sludge phylotypes (bold typeface) and isolated members of the phylum Chloroflexi. The alignment used for the tree applied a 20% conservational filter to remove hypervariable positions, giving 1108 aligned positions. Phylogenetic classification is indicated with black brackets and is based on the MiDAS database (Release 2.1). Probe coverage of probes relevant to the current study is shown with red coloured brackets. Associated activated sludge morphotypes of previously described phylotypes are designated with blue coloured brackets. Bootstrap values from 100 resamplings are indicated for branches with >50% (white dot), 50-70% (gray), and >90% (black) support. Species of the phylum Cyanobacteria were used as the outgroup. The scale bar represents substitutions per nucleotide base.

### Phylogeny and FISH probe design

*‘Ca*. Villigracilis’ are members of the Anaerolineaceae, which is currently the sole family of the class Anaerolineae in the MiDAS and SILVA taxonomies. These sequences fall within order SBR1031 in the Greengenes taxonomy (McDonald *et al*. 2012). The CFX763A and CFX763B probes were designed to cover separate sub-groups (A and B) of the *‘Ca*. Villigracilis’ (**Figure S2**) - collectively covering >60% of the MiDAS database sequences classified to the genus. The target region is not covered by the V1-3 region amplicon sequences, although both probes match the full-length database sequences most closely related to the abundant OTU sequences (data not shown). When applied to full-scale activated sludge biomass, both probes hybridized thin filaments (0.3-0.4 μm wide and 15-50 μm long) that were often observed in bundles and almost exclusively located within the flocs. The CFX763AB_H1A and CFX763AB_H1B helper probes are recommended to give optimal fluorescence signal for both probes. Amplicon sequencing of the V1-3 region of the 16S rRNA cannot be used to confidently separate the A and B sub-groups, due to high sequence similarity, but qFISH indicates that the former is the more numerically important of the two.

The *‘Ca*. Amarolinea’ genus falls within the novel MiDAS Chloroflexi class-level-group SJA-15, together with the also abundant genus *‘Ca*. Sarcinithrix’ (**Figure 2**). A probe to cover the entire *‘Ca*. Amarolinea’ group was not identified, so the CFX64 (**Table 1**) was designed for the abundant amplicon OTUs (OTU_3 and OTU_4592, **Figure S3**) and the most closely related full-length database sequence (AF513086). These sequences share >97% similarity. As the probe covers these abundant OTUs, it should cover the majority of members of the genus in full-scale activated sludge in Denmark. When applied to activated sludge, the probe hybridized exclusively to filamentous bacteria (see later for a detailed description of their morphology). A strong positive signal was obtained without addition of designed unlabeled helper probes CFX64_H1 and CFX64_H2 (**Table 1**), which did not noticeably improve fluorescent signal (data not shown). The *‘Ca*. Amarolinea’ filaments constituted up to 30% of the community biovolume in some full-scale WWTPs in Denmark - confirming their high abundance with amplicon sequencing (see **Table 2**).

**Table 2.** Abundance estimation: 16S rRNA amplicon sequencing and qFISH (percentage of total).

| WWTP | Sample date | Abundance (%) |  |
| --- | --- | --- | --- |
|  |  | Sequencing* | qFISH |
| **‘Ca. Villigracilis’** |  |  |  |
| Odense North East | September 2015 | 9.0 | sub-group A: 12 ± 4<br>sub-group B: 1 ± 1 |
| Aalborg East | August 2015 | 6.0 | sub-group A: 11 ± 3<br>sub-group B: > 1 |
| Horsens | August 2006 | 6.0 | sub-group A: 11 ± 2<br>sub-group B: > 1 |
| Fredericia | October 2015 | 3.3 | sub-group A: 6 ± 1<br>sub-group B: > 1 |
| **‘Ca. Amarolinea’** |  |  |  |
| Aalborg West | May 2014 | 11.9 | 15.8 |
| Bjergmarken | August 2013 | 33.7 | 30.2 |
| Odense North West | August 2010 | 24.2 | 16.8 |
| **‘Ca. Sarcinithrix’** |  |  |  |
| Viby | August 2008 | 2.3 | 2 |
| Boeslum | August 2008 | 2.1 | 3 |
| Bjergmarken | February 2013 | 1.2 | 2 |
| Ejby Mølle | May 2008 | 1.0 | 1 |
| Aalborg West | February 2012 | 0.8 | 1 |

\* Taken from the 16S rRNA gene amplicon sequencing MiDAS survey of Danish WWTPs (McIlroy *et al.* 2015).

**Figure 2.**
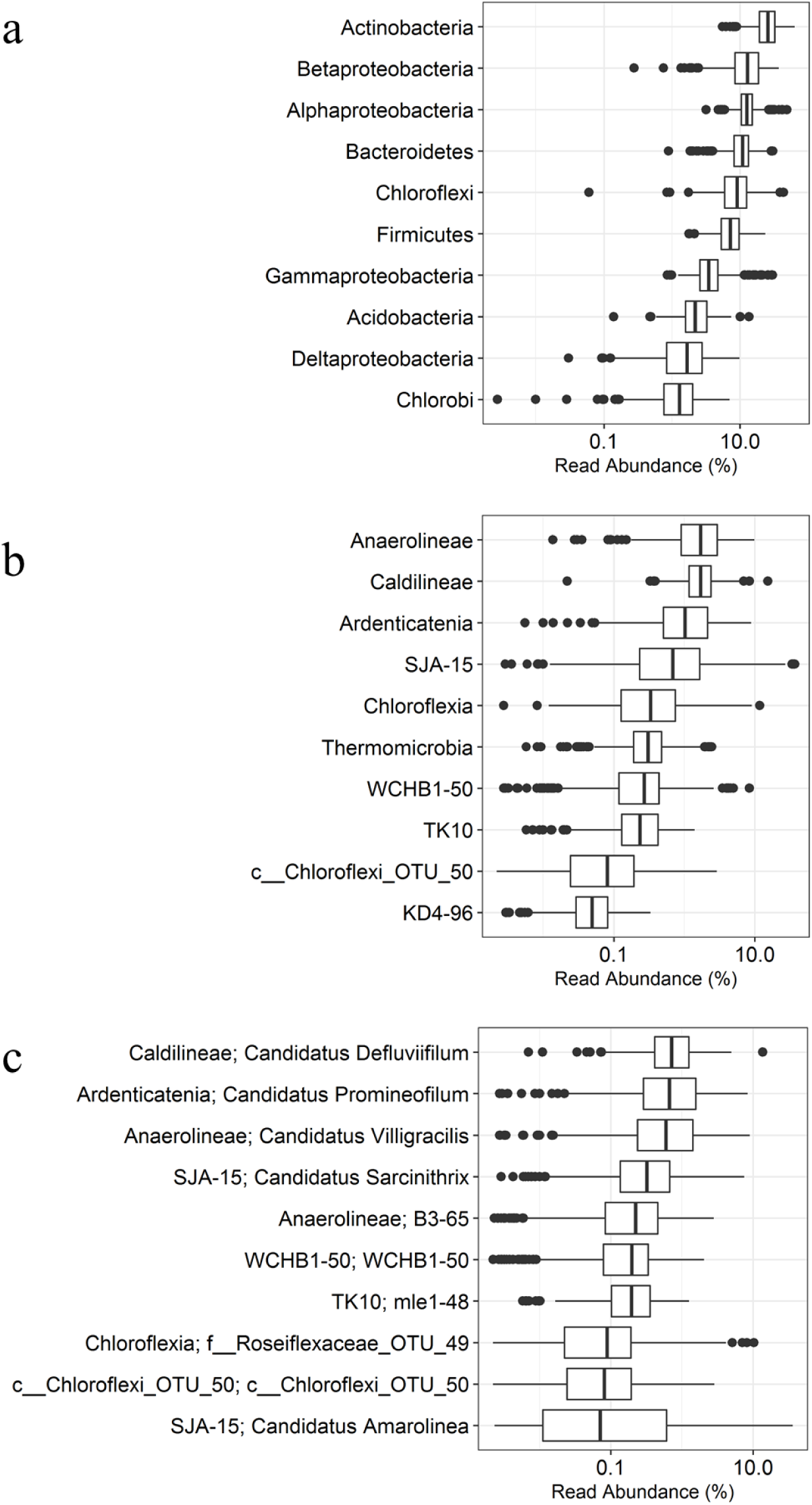
Distribution of Chloroflexi in 25 full-scale activated sludge WWTPs sampled 2-4 times per year from 2006 to 2015. (a) 10 most abundant phyla in Danish WWTPs. (b) 10 most abundant Chloroflexi classes in Danish WWTPs (c) 10 most abundant Chloroflexi genera in Danish WWTPs. X-axis shows the relative read abundance in percentage of total bacteria.

**Figure 3.**
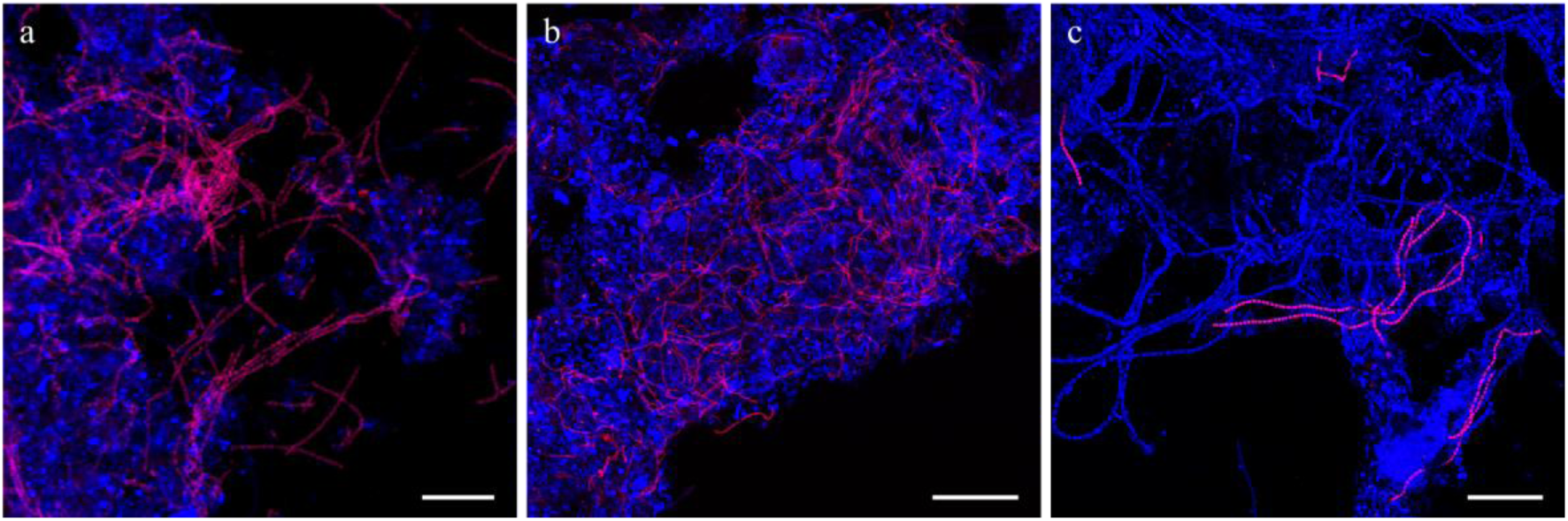
Composite FISH micrographs of novel Chloroflexi genera in full-scale activated sludge. Specific probes (Cy3-label, red) target (a) *‘Ca*. Amarolinea’, (b) *‘Ca*. Villigracilis’, and (c) *‘Ca*. Sarcinithrix’, and EUBmix probe (Cy5-label, blue) targets most bacteria. Activated sludge was sampled from (a) Bjergmarken WWTP, (b) Odense North East WWTP, and (c) Aalborg West WWTP. Target filaments appear magenta, while all other cells appear blue. Scale bars represent 20 μm.

**Figure 4.**
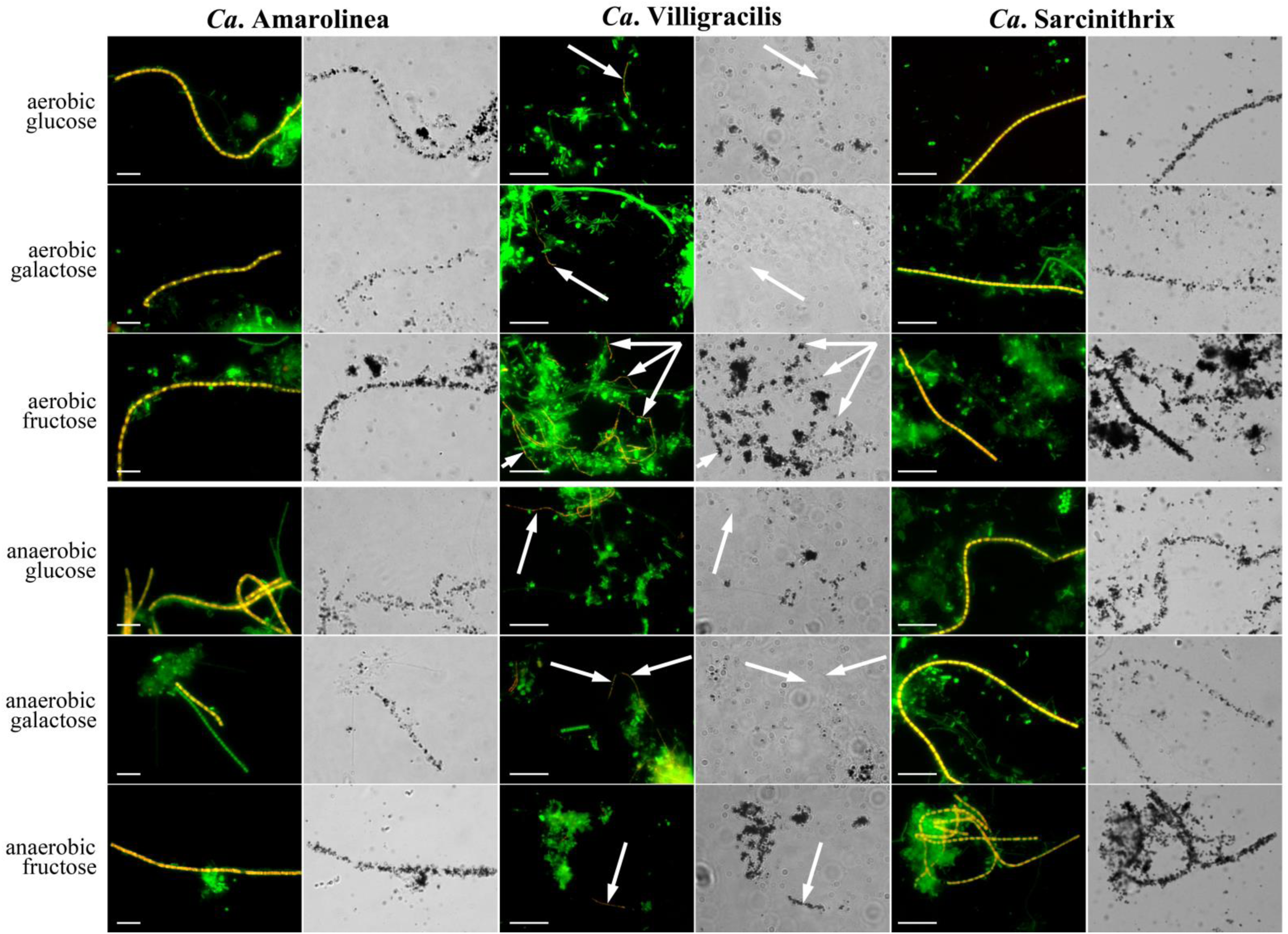
FISH and corresponding bright-field MAR micrographs showing sugar uptake by three abundant Chloroflexi phylotypes (‘*Ca*. Amarolinea’ hybridized probe CFX64; *‘Ca*. Villigracilis’ sub-group A hybridized probe CFX763A; *‘Ca*. Sarcinithrix’ hybridized probe CFX1151). Activated sludge was sampled in Aalborg West or Odense North West WWTPs. Target cells in FISH micrograph overlays appear yellow: specific probe (red) + EUBmix (green); and non-target cells appear green (EUBmix only). Black silver granules indicate positive MAR signal. Scale bar represents 10 μm.

FISH probes are already available to target the *‘Ca*. Sarcinithrix’ (CFX67), which was shown to possess the Eikelboom 0914 morphotype in nutrient removal activated sludge systems in Australia (Speirs *et al*. 2011). Application of CFX67 probe did not give significant positive fluorescence to bacterial cells in Danish plants. The probe misses several full-length sequences classified to the genus and does not cover any of the abundant OTU sequences (>0.1% average read abundance in at least 1 plant) from the MiDAS full-scale survey. As such, two new probes were designed to give better coverage of the clade. The designed CFX449 and CFX1151 individually target 85% of the full-length database sequences. As such, they can be applied with different fluorochromes to confirm specific coverage of the genus, or together as a mix to give higher signal (**Figure S4**). Helper probes were not required for either probe, but did give a more even signal over the filament, which was also reported by Speirs *et al*. (2011) for the CFX67 probe. The few filaments positive for the CFX67 probe in the Danish WWTPs assessed in this study also hybridized the CFX449 and CFX1151 probes (**Figure S4**). Quantitative FISH values with the CFX449 and CFX1151 were similar to amplicon-sequencing based estimates (**Table 2**).

All three phylotypes studied gave positive hybridization signal with the EUBmix probe set, which is commonly applied as a universal probe targeting bacteria (EUB338, EUB338-II, and EUB338-III). This is of interest, given that many Chloroflexi reportedly lack the target site for the EUBmix probe set and fail to hybridize the probe *in situ* (Kragelund *et al*. 2007, 2011; Speirs *et al*. 2009). Most 16S rRNA gene sequences of the *‘Ca*. Amarolinea’, *‘Ca*. Sarcinithrix’ and ‘Kouleothrix’ contain the site for the EUB338 probe, the *‘Ca*. Villigracilis’ sequences possess the EUB338-III site, and all groups have been shown to hybridize the probe *in situ*. Most members of the ‘*Ca*. Defluviifilum’ have one mismatch, though *in silico* analysis with the MathFISH software (Yilmaz, Parnerkar and Noguera 2011) predicts positive binding (with a calculated melting formamide point ([ΔFA]_m_) of 50%), which is confirmed *in situ* for filaments hybridizing the T0803-0654 probe designed to target the group (Kragelund *et al*. 2011). Thus, of the abundant phylotypes, only the *‘Ca*. Promineofilum’ genus is not covered by the EUBmix probe set (Speirs *et al*. 2009).

### Morphological description and classification

The morphological properties of the CFX64 positive filaments, representing the *‘Ca*. Amarolinea’ genus, were investigated in detail for association to a morphotype of the well-known antecedent classification systems (Eikelboom 1975). Though not clearly visible, the cells appeared to be rectangular with no visible septa, a trichrome thickness of 1.0-2.2 μm, and a length in the 20-140 μm range. They were non-motile, Gram stain negative, with no branching or attached growth. The whole filaments stained blue/violet with the Neisser stain, with no visible volutin granules. Excess polyphosphate stores were not observed with DAPI staining. Cells did not appear to contain excess stores of polyhydroxyalkanoates (PHAs), with negative results with Nile blue A staining and only small positive granules observed with the Sudan black stain. From these observations, primarily based on the characteristic violet color of the cells after Neisser staining, it is suggested that the morphology of the filament is most consistent with the Eikelboom type 0092 morphotype (Eikelboom 2000). The *‘Ca*. Promineofilum’ also reportedly has the 0092 morphotype (Speirs *et al*. 2009), but there was no observed overlap between the CFX64 and the CFX197 probes targeting the ‘*Ca*. Amarilimum’ and ‘*Ca*. Promineofilum’ genera, respectively (**Figure S5**). The *‘Ca*. Promineofilum’ phylotype is also thinner in appearance, with a trichome diameter of approx. 0.8 μm (Speirs *et al*. 2009). Very few of the fluorescent sulphate modified microspheres attached to the CFX64 positive filaments (data not shown), indicating that they do not have a hydrophobic surface and are likely not involved in foam formation.

Morphological descriptions are already reported for members of the *‘Ca*. Sarcinithrix’ (Speirs *et al*. 2011) and *‘Ca*. Defluviifilum’ (Kragelund *et al*. 2011; Speirs *et al*. 2017). The surface hydrophobicity of *‘Ca*. Sarcinithrix’ was assessed for the first time here, where it was determined to be hydrophilic and therefore unlikely to be involved in foam formation. Morphological classification of the *‘Ca*. Villigracilis’ was not successful due to their location within the floc, making interpretation of staining analyses difficult.

### In situ substrate uptake

The results for substrate uptake by probe-defined *‘Ca*. Villigracilis’ sub-groups A and B, *‘Ca*. Amarolinea’, and *‘Ca*. Sarcinithrix’ using MAR-FISH under various conditions are shown in **Table S1**, and a summary of known *in situ* traits of abundant Chloroflexi is given in **Table 3**. ‘*Ca*. Villigracilis’ sub-group A, ‘*Ca*. Amarolinea’, and *‘Ca*. Sarcinithrix’ only utilized sugars of the 13 substrates tested, consistent with other characterized Chloroflexi genera. The phylotypes differed in the types of sugars taken up, noting that variation was also observed within the *‘Ca*. Amarolinea’ genus – with some filaments strongly positive and others clearly negative for fructose uptake. *‘Ca*. Villigracilis’ sub-group B filaments were negative for all substrates and conditions. It may be that they have a relatively lower activity than the much more abundant sub-group A filaments that is below the detection of MAR. Further analyses are required to assess the reason for the observed lack of substrate uptake. All three genera were able to take up substrates under anoxic conditions, suggesting fermentative metabolisms. Anoxic uptake of sugars in presence of nitrate/nitrite was also observed, but their potential for denitrification is unclear, given that uptake was also observed without nitrate/nitrite addition. The same ambiguous results were obtained for the *‘Ca*. Defluviifilum’ (Kragelund *et al*. 2011). The ability for nitrification was also assessed for these groups in this study, given that *Nitrolancetus hollandicus*, a nitrite oxidizing member of the class Thermomicrobia of the Chloroflexi, was isolated from activated sludge (Sorokin *et al*. 2012); albeit in a different class to the abundant members of the Chloroflexi Danish full-scale systems. None of the genera appeared to be behaving as nitrifiers, with no observed uptake of labeled CO_2_ in the presence of ammonia (**Table 3 and S1**), while a positive MAR signal was noted for *Nitrosomonas* as well as *Nitrospira*, targeted by the probe Cluster6a_192 (Adamczyk *et al*. 2003) and Ntsp662 (Daims *et al*. 2001), respectively.

**Table 3.**
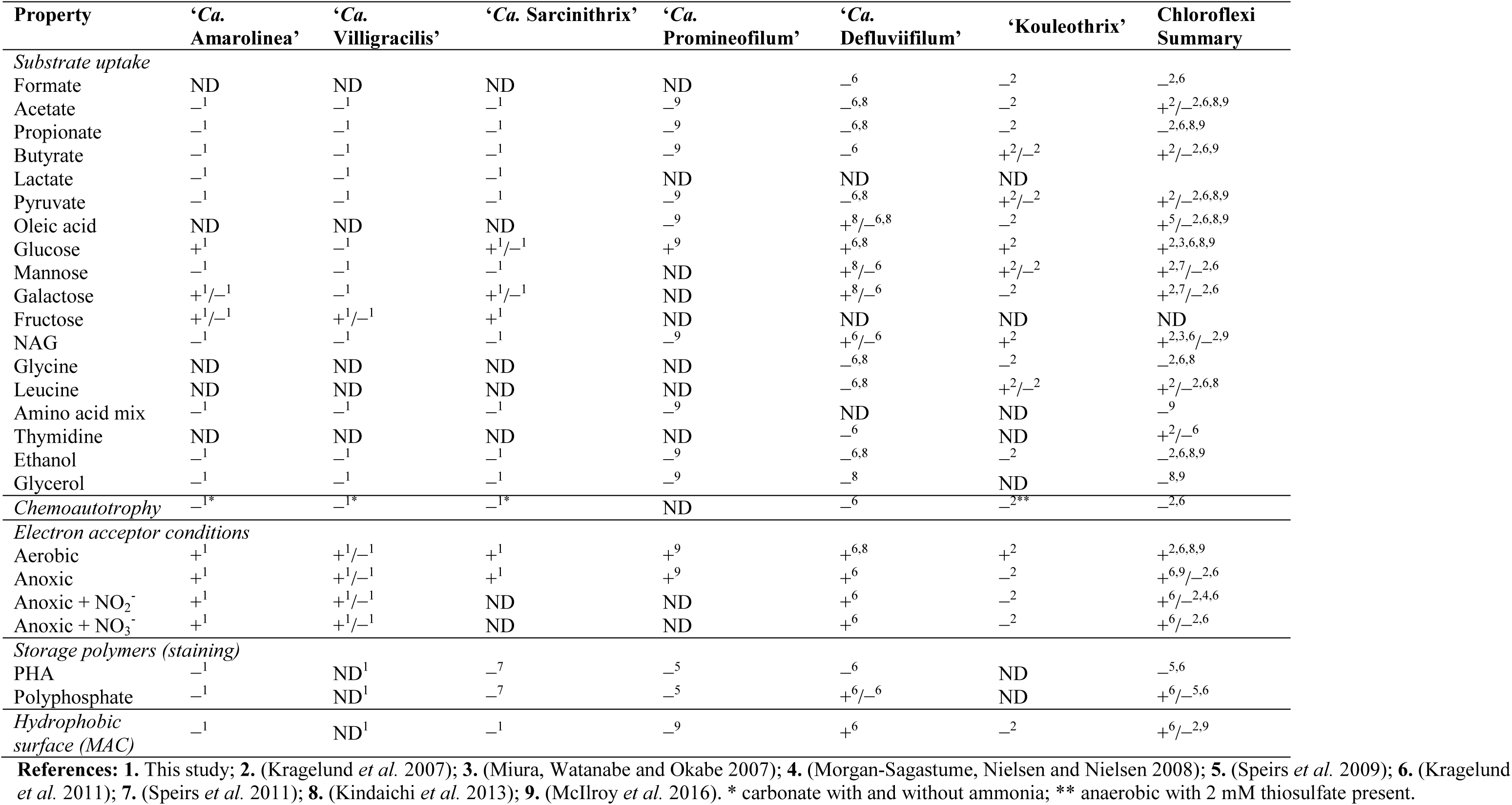
Summary of the known *in situ* physiology for Chloroflexi commonly found in activated sludge.

| Property | 'Ca. Amarolinea' | 'Ca. Villigracilis' | 'Ca. Sarcinithrix' | 'Ca. Promineofilum' | 'Ca. Defluviifilum' | 'Kouleothrix' | Chloroflexi Summary |
| --- | --- | --- | --- | --- | --- | --- | --- |
| *Substrate uptake* |  |  |  |  |  |  |  |
| Formate | ND | ND | ND | ND | -<sup>6</sup> | -<sup>2</sup> | -<sup>2,6</sup> |
| Acetate | -<sup>1</sup> | -<sup>1</sup> | -<sup>1</sup> | -<sup>9</sup> | -<sup>6,8</sup> | -<sup>2</sup> | +<sup>2</sup>/<sub>-</sub><sup>2,6,8,9</sup> |
| Propionate | -<sup>1</sup> | -<sup>1</sup> | -<sup>1</sup> | -<sup>9</sup> | -<sup>6,8</sup> | -<sup>2</sup> | -<sup>2,6,8,9</sup> |
| Butyrate | -<sup>1</sup> | -<sup>1</sup> | -<sup>1</sup> | -<sup>9</sup> | -<sup>6</sup> | +<sup>2</sup>/<sub>-</sub><sup>2</sup> | +<sup>2</sup>/<sub>-</sub><sup>2,6,9</sup> |
| Lactate | -<sup>1</sup> | -<sup>1</sup> | -<sup>1</sup> | ND | ND | ND |  |
| Pyruvate | -<sup>1</sup> | -<sup>1</sup> | -<sup>1</sup> | -<sup>9</sup> | -<sup>6,8</sup> | +<sup>2</sup>/<sub>-</sub><sup>2</sup> | +<sup>2</sup>/<sub>-</sub><sup>2,6,8,9</sup> |
| Oleic acid | ND | ND | ND | -<sup>9</sup> | +<sup>8</sup>/<sub>-</sub><sup>6,8</sup> | -<sup>2</sup> | +<sup>5</sup>/<sub>-</sub><sup>2,6,8,9</sup> |
| Glucose | +<sup>1</sup> | -<sup>1</sup> | +<sup>1</sup>/<sub>-</sub><sup>1</sup> | +<sup>9</sup> | +<sup>6,8</sup> | +<sup>2</sup> | +<sup>2,3,6,8,9</sup> |
| Mannose | -<sup>1</sup> | -<sup>1</sup> | -<sup>1</sup> | ND | +<sup>8</sup>/<sub>-</sub><sup>6</sup> | +<sup>2</sup>/<sub>-</sub><sup>2</sup> | +<sup>2,7</sup>/<sub>-</sub><sup>2,6</sup> |
| Galactose | +<sup>1</sup>/<sub>-</sub><sup>1</sup> | -<sup>1</sup> | +<sup>1</sup>/<sub>-</sub><sup>1</sup> | ND | +<sup>8</sup>/<sub>-</sub><sup>6</sup> | -<sup>2</sup> | +<sup>2,7</sup>/<sub>-</sub><sup>2,6</sup> |
| Fructose | +<sup>1</sup>/<sub>-</sub><sup>1</sup> | +<sup>1</sup>/<sub>-</sub><sup>1</sup> | +<sup>1</sup> | ND | ND | ND | ND |
| NAG | -<sup>1</sup> | -<sup>1</sup> | -<sup>1</sup> | -<sup>9</sup> | +<sup>6</sup>/<sub>-</sub><sup>6</sup> | +<sup>2</sup> | +<sup>2,3,6</sup>/<sub>-</sub><sup>2,9</sup> |
| Glycine | ND | ND | ND | ND | -<sup>6,8</sup> | -<sup>2</sup> | -<sup>2,6,8</sup> |
| Leucine | ND | ND | ND | ND | -<sup>6,8</sup> | +<sup>2</sup>/<sub>-</sub><sup>2</sup> | +<sup>2</sup>/<sub>-</sub><sup>2,6,8</sup> |
| Amino acid mix | -<sup>1</sup> | -<sup>1</sup> | -<sup>1</sup> | -<sup>9</sup> | ND | ND | -<sup>9</sup> |
| Thymidine | ND | ND | ND | ND | -<sup>6</sup> | ND | +<sup>2</sup>/<sub>-</sub><sup>6</sup> |
| Ethanol | -<sup>1</sup> | -<sup>1</sup> | -<sup>1</sup> | -<sup>9</sup> | -<sup>6,8</sup> | -<sup>2</sup> | -<sup>2,6,8,9</sup> |
| Glycerol | -<sup>1</sup> | -<sup>1</sup> | -<sup>1</sup> | -<sup>9</sup> | -<sup>8</sup> | ND | -<sup>8,9</sup> |
| *Chemoautotrophy* | -<sup>1</sup>* | -<sup>1</sup>* | -<sup>1</sup>* | ND | -<sup>6</sup> | -<sup>2</sup>** | -<sup>2,6</sup> |
| *Electron acceptor conditions* |  |  |  |  |  |  |  |
| Aerobic | +<sup>1</sup> | +<sup>1</sup>/<sub>-</sub><sup>1</sup> | +<sup>1</sup> | +<sup>9</sup> | +<sup>6,8</sup> | +<sup>2</sup> | +<sup>2,6,8,9</sup> |
| Anoxic | +<sup>1</sup> | +<sup>1</sup>/<sub>-</sub><sup>1</sup> | +<sup>1</sup> | +<sup>9</sup> | +<sup>6</sup> | -<sup>2</sup> | +<sup>6,9</sup>/<sub>-</sub><sup>2,6</sup> |
| Anoxic + NO<sub>2</sub><sup>-</sup> | +<sup>1</sup> | +<sup>1</sup>/<sub>-</sub><sup>1</sup> | ND | ND | +<sup>6</sup> | -<sup>2</sup> | +<sup>6</sup>/<sub>-</sub><sup>2,4,6</sup> |
| Anoxic + NO<sub>3</sub><sup>-</sup> | +<sup>1</sup> | +<sup>1</sup>/<sub>-</sub><sup>1</sup> | ND | ND | +<sup>6</sup> | -<sup>2</sup> | +<sup>6</sup>/<sub>-</sub><sup>2,6</sup> |
| *Storage polymers (staining)* |  |  |  |  |  |  |  |
| PHA | -<sup>1</sup> | ND<sup>1</sup> | -<sup>7</sup> | -<sup>5</sup> | -<sup>6</sup> | ND | -<sup>5,6</sup> |
| Polyphosphate | -<sup>1</sup> | ND<sup>1</sup> | -<sup>7</sup> | -<sup>5</sup> | +<sup>6</sup>/<sub>-</sub><sup>6</sup> | ND | +<sup>6</sup>/<sub>-</sub><sup>5,6</sup> |
| *Hydrophobic surface (MAC)* | -<sup>1</sup> | ND<sup>1</sup> | -<sup>1</sup> | -<sup>9</sup> | +<sup>6</sup> | -<sup>2</sup> | +<sup>6</sup>/<sub>-</sub><sup>2,9</sup> |

**References:** **1.** This study; **2.** (Kragelund *et al.* 2007); **3.** (Miura, Watanabe and Okabe 2007); **4.** (Morgan-Sagastume, Nielsen and Nielsen 2008); **5.** (Speirs *et al.* 2009); **6.** (Kragelund *et al.* 2011); **7.** (Speirs *et al.* 2011); **8.** (Kindaichi *et al.* 2013); **9.** (McIlroy *et al.* 2016). \* carbonate with and without ammonia; \*\* anaerobic with 2 mM thiosulfate present.

## Discussion

The results of this study of the distribution and *in situ* morphology and physiology of individual abundant phylotypes of the phylum Chloroflexi give valuable insight into their potential importance to the operation of WWTPs. The *‘Ca*. Promineofilum’, *‘Ca*. Defluviifilum’, *‘Ca*. Villigracilis’, and *‘Ca*. Sarcinithrix’ appear to be consistently abundant filamentous members of the full-scale WWTP community, where they certainly make an important contribution to the bulk nutrient transformations. Relative to other abundant Chloroflexi genera, the distribution of the *‘Ca*. Amarolinea’ is much more dynamic, reaching levels of >30% of the biovolume in some plants. As such, it is more likely that the *‘Ca*. Amarolinea’ are responsible for acute bulking episodes in Danish WWTPs, while other abundant phylotypes are core members of the community that are possibly important to floc structure and the breakdown of organics (see later). While the *‘Ca*. Villigracilis’ are filamentous, they are almost exclusively found within the flocs and are unlikely to contribute to bulking episodes. This observation explains why they have never been detected before and highlights the value of large-scale full-scale surveys, such as MiDAS, for describing the abundant members of wastewater treatment systems and the importance of *in situ* analyses to evaluate the role of organisms in floc structure and settleability. In addition to a potential contribution to bulking, members of the *‘Ca*. Promineofilum’ are also putatively involved in membrane fouling in MBR systems (Ziegler *et al*. 2016), and *‘Ca*. Defluviifilum’ has a hydrophobic cell surface that has been implicated in the stabilization of problematic foams on the surface of reactor tanks (Kragelund *et al*. 2011).

In recent studies Speirs *et al*., (2015; 2017) described filamentous Chloroflexi phylotypes in Australian WWTPs possessing the bulking Eikelboom morphotypes 0803 and 0041/0675 that are commonly observed by light microscopy in Danish plants (Mielczarek *et al*. 2012). These phylotypes classified within the Anaerlolineae and Caldilinaea, respectively. However, none of these were found to be abundant in the amplicon based survey of Danish systems (data not shown). Their FISH probe defined 0675 phylotype falls within the MiDAS defined *Ca*. Defluviifilum’ genus (v. 2.1), shown previously to possess the Eikelboom 0803 phylotype (Kragelund *et al*. 2011), which they suggest should be split into a separate genus based on the divergence of its 16S rRNA gene sequence (only *Ca*. 90% similar). Further characterisation of members of the current *‘Ca*. Defluviifilum’ clade, including obtaining representative genomes, will help to resolve the phylogeny of these organisms. Members of the ‘Kouleothrix’ genus, shown to possess the Eikelboom 1851 bulking morphotype (Beer *et al*. 2002; Kohno, Sei and Mori 2002), were also relatively low in abundance in Danish plants (Kragelund *et al*. 2011) but are common in other countries – such as Japan, where they have been suggested as a major contributor to filamentous bulking episodes (Nittami *et al*. 2017). Global surveys will help to establish how relevant the abundant Danish phylotypes characterised in this paper are worldwide.

The abundant phylotypes in Danish WWTPs appear to be organoheterotrophic, fermentative, facultative anaerobes. This is suggested from their exclusive utilization of sugars and ability for anaerobic carbon uptake. Fermentative pathways were also annotated in the sole genome from these genera – *Ca*. Promineofilum breve’ (McIlroy *et al*. 2016). The physiology determined for the *‘Ca*. Defluviifilum’ and *Ca*. Villigracilis’ is consistent with other members of their respective classes (Caldilineae and Anaerolinea), which are mostly fermentative organoheterotrophic filaments growing on sugars and/or amino acids (Sekiguchi *et al*. 2003; Yamada *et al*. 2006, 2007; Grégoire *et al*. 2011; Nunoura *et al*. 2013; Podosokorskaya *et al*. 2013; Imachi *et al*. 2014; McIlroy *et al*. 2017a). *‘Ca*. Villigracilis’ is the first reported facultative anaerobic genus of the Anaerolineae, with all isolates of other described genera being obligate anaerobes.

In addition to the likely fermentation of sugars by the abundant Chloroflexi, metabolic diversity within the less abundant members of the phylum is evident – with reported assimilation of short and long chain fatty acids, amino acids, and glycerol (**Table 3**). Some members of the phylum also appear unable to take up substrates anaerobically and may scavenge sugars released from aerobic breakdown of complex organic matter. Uptake of *N*-aminoglucosamine (NAG), a component of peptidoglycan and lipopolysaccharides, for some Chloroflexi, suggests a specific role in the breakdown of cellular material (Kragelund *et al*. 2007, 2011; Miura, Watanabe and Okabe 2007). Of the known Chloroflexi genera reportedly abundant in activated sludge, only the ‘Kouleothrix’ genus is seemingly unable to take up substrates anaerobically *in situ* (Kragelund *et al*. 2007), although isolates of the genus are capable of anaerobic fermentative growth on sugars (Kohno, Sei and Mori 2002).

In this study, the most abundant members of the Chloroflexi in Danish nutrient removal WWTPs were identified and their ecophysiology described. These phylotypes appear to differ in their impact on plant operation - with suggested importance in sludge settleability, foaming, and membrane fouling being associated with different groups. All abundant members of the phylum likely ferment sugars, and future research should aim to obtain representative genomes for each in order to carry out more detailed comparison of their metabolic activities. Such an approach will explain important questions regarding how these organisms coexist, and what conditions determine their relative abundances. The FISH probes designed in this study will allow more hypothesis-based *in situ* investigation of their physiologies, based on genomic evidence. The taxonomic annotation and design of FISH probes for the abundant Chloroflexi, in combination with the high throughput nature of 16S rRNA gene amplicon sequencing, will also allow their routine observation and study. Defining and naming these novel genus level taxa importantly provides the foundation, upon which information on their morphology, distribution, and physiology can be gathered for an in-depth understanding of their ecology and how this might relate to operational parameters.

## Acknowledgements

We thank K. Vilstrup for help to latin names.

## References

Adamczyk J, Hesselsoe M, Iversen N et al. The Isotope Array, a New Tool That Employs Substrate-Mediated Labeling of rRNA for Determination of Microbial Community Structure and Function. Appl Environ Microbiol 2003;69:6875–87.

Amann RI, Binder BJ, Olson RJ et al. Combination of 16S rRNA-targeted oligonucleotide probes with flow cytometry for analyzing mixed microbial populations. Appl Env Microbiol 1990;56:1919–25.

Beer M, Seviour EM, Kong Y et al. Phylogeny of the filamentous bacterium Eikelboom Type 1851, and design and application of a 16S rRNA targeted oligonucleotide probe for its fluorescence in situ identification in activated sludge. FEMS Microbiol Lett 2002;207:179–83.

Beer M, Stratton HM, Griffiths PC et al. Which are the polyphosphate accumulating organisms in full-scale activated sludge enhanced biological phosphate removal systems in Australia? J Appl Microbiol 2006;100:223–43.

Björnsson L, Hugenholtz P, Tyson GW et al. Filamentous Chloroflexi (green non-sulfur bacteria) are abundant in wastewater treatment processes with biological nutrient removal. Microbiology 2002;148:2309–18.

Cole JR, Wang Q, Fish JA et al. Ribosomal Database Project: data and tools for high throughput rRNA analysis. Nucleic Acids Res 2014;42:D633–42.

Daims H, Brühl A, Amann R et al. The domain-specific probe EUB338 is insufficient for the detection of all Bacteria: development and evaluation of a more comprehensive probe set. Syst Appl Microbiol 1999;22:434–44.

Daims H, Lücker S, Wagner M. daime, a novel image analysis program for microbial ecology and biofilm research. Env Microbiol 2006;8:200–13.

Daims H, Nielsen JL, Nielsen PH et al. In Situ Characterization of Nitrospira-Like Nitrite-Oxidizing Bacteria Active in Wastewater Treatment Plants. Appl Environ Microbiol 2001;67:5273–84.

Daims H, Stoecker K, Wagner M. Fluorescence in situ hybridization for the detection of prokaryotes. In: Osborn AM, Smith CJ (eds.). Molecular Microbial Ecology. New York: Taylor & Francis, 2005, 213–39.

Eikelboom D. Process Control of Activated Sludge Plants by Microscopic Investigation. London: IWA Publishing, 2000.

Eikelboom DH. Filamentous organisms observed in activated sludge. Water Res 1975;9:365–88.

Gich F, Garcia-Gil J, Overmann J. Previously unknown and phylogenetically diverse members of the green nonsulfur bacteria are indigenous to freshwater lakes. Arch Microbiol 2001;177:1–10.

Grégoire P, Bohli M, Cayol J-L et al. Caldilinea tarbellica sp. nov., a filamentous, thermophilic, anaerobic bacterium isolated from a deep hot aquifer in the Aquitaine Basin. Int J Syst Evol Microbiol 2011;61:1436–41.

Greuter D, Loy A, Horn M et al. probeBase—an online resource for rRNA-targeted oligonucleotide probes and primers: new features 2016. Nucleic Acids Res 2016;44:D586–9.

Imachi H, Sakai S, Lipp JS et al. Pelolinea submarina gen. nov., sp. nov., an anaerobic, filamentous bacterium of the phylum Chloroflexi isolated from subseafloor sediment. Int J Syst Evol Microbiol 2014;64:812–8.

Jenkins D, Richard MG, Daigger GT. Manual on the Causes and Control of Activated Sludge Bulking, Foaming and Other Solids Separation Problems. 3 rd. London, England: CRC Press LLC, 2004.

Kindaichi T, Nierychlo M, Kragelund C et al. High and stable substrate specificities of microorganisms in enhanced biological phosphorus removal plants. Environ Microbiol 2013;15:1821–31.

Kohno T, Sei K, Mori K. Characterization of type 1851 organism isolated from activated sludge samples. Water Sci Technol J Int Assoc Water Pollut Res 2002;46:111–4.

Kragelund C, Levantesi C, Borger A et al. Identity, abundance and ecophysiology of filamentous Chloroflexi species present in activated sludge treatment plants. FEMS Microbiol Ecol 2007;59:671–82.

Kragelund C, Müller E, Schade M et al. Identification of filamentous bacteria by FISH. In: Nielsen PH, Daims H, Lemmer H (eds.). FISH Handbook for Biological Wastewater Treatment. London: IWA Publishing, 2009, 33–68.

Kragelund C, Nielsen JL, Thomsen TR et al. Ecophysiology of the filamentous Alphaproteobacterium Meganema perideroedes in activated sludge. FEMS Microbiol Ecol 2005;54:111–22.

Kragelund C, Thomsen TR, Mielczarek AT et al. Eikelboom’s morphotype 0803 in activated sludge belongs to the genus Caldilinea in the phylum Chloroflexi. FEMS Microbiol Ecol 2011;76:451–62.

Ludwig W, Strunk O, Westram R et al. ARB: a software environment for sequence data. Nucleic Acids Res 2004;32:1363–71.

McDonald D, Price MN, Goodrich J et al. An improved Greengenes taxonomy with explicit ranks for ecological and evolutionary analyses of bacteria and archaea. ISME J 2012;6:610–8.

McIlroy SJ, Karst SM, Nierychlo M et al. Genomic and in situ investigations of the novel uncultured Chloroflexi associated with 0092 morphotype filamentous bulking in activated sludge. ISME J 2016;In press.

McIlroy SJ, Kirkegaard RH, Dueholm MS et al. Culture-Independent Analyses Reveal Novel Anaerolineaceae as Abundant Primary Fermenters in Anaerobic Digesters Treating Waste Activated Sludge. Front Microbiol 2017a;8, DOI: 10.3389/fmicb.2017.01134.

McIlroy SJ, Kirkegaard RH, McIlroy B et al. MiDAS 2.0: an ecosystem-specific taxonomy and online database for the organisms of wastewater treatment systems expanded for anaerobic digester groups. Database 2017b;2017, DOI: 10.1093/database/bax016.

McIlroy SJ, Saunders AM, Albertsen M et al. MiDAS: the field guide to the microbes of activated sludge. Database 2015;2015:1–8.

McIlroy SJ, Tillett D, Petrovski S et al. Non-target sites with single nucleotide insertions or deletions are frequently found in 16S rRNA sequences and can lead to false positives in fluorescence in situ hybridization (FISH). Env Microbiol 2011;13:38–47.

Mielczarek AT, Kragelund C, Eriksen PS et al. Population dynamics of filamentous bacteria in Danish wastewater treatment plants with nutrient removal. Water Res 2012;46:3781–95.

Mielczarek AT, Nguyen HT, Nielsen JL et al. Population dynamics of bacteria involved in enhanced biological phosphorus removal in Danish wastewater treatment plants. Water Res 2013;47:1529–44.

Miura Y, Watanabe Y, Okabe S. Significance of Chloroflexi in performance of submerged membrane bioreactors (MBR) treating municipal wastewater. Environ Sci Technol 2007;41:7787–94.

Morgan-Sagastume F, Nielsen JL, Nielsen PH. Substrate-dependent denitrification of abundant probe-defined denitrifying bacteria in activated sludge. FEMS Microbiol Ecol 2008;66:447–61.

Murray RG, Stackebrandt E. Taxonomic note: implementation of the provisional status Candidatus for incompletely described procaryotes. Int J Syst Bacteriol 1995;45:186–7.

Nierychlo M, Nielsen JL, Nielsen PH. Studies of the Ecophysiology of Single Cells in Microbial Communities by (Quantitative) Microautoradiography and Fluorescence In Situ Hybridization (MAR-FISH). In: McGenity TJ, Timmis KN, Nogales Fernández B (eds.). Hydrocarbon and Lipid Microbiology Protocols, Springer Protocols Handbooks. 1st ed. Berlin-Heidelberg: Springer-Verlag, 2015.

Nittami T, Speirs LBM, Yamada T et al. Quantification of *Chloroflexi* Eikelboom morphotype 1851 for prediction and control of bulking events in municipal activated sludge plants in Japan. Appl Microbiol Biotechnol 2017;101:3861–9.

Nunoura T, Hirai M, Miyazaki M et al. Isolation and characterization of a thermophilic, obligately anaerobic and heterotrophic marine Chloroflexi bacterium from a Chloroflexi dominated microbial community associated with a japanese shallow hydrothermal system, and proposal for Thermomarinilin. Microbes Environ JSME 2013;28:228–35.

Ostle AG, Holt JG. Nile blue A as a fluorescent stain for poly-beta-hydroxybutyrate. Appl Env Microbiol 1982;44:238–41.

Podosokorskaya OA, Bonch-Osmolovskaya EA, Novikov AA et al. Ornatilinea apprima gen. nov., sp. nov., a cellulolytic representative of the class Anaerolineae. Int J Syst Evol Microbiol 2013;63:86–92.

Quast C, Pruesse E, Yilmaz P et al. The SILVA ribosomal RNA gene database project: improved data processing and web-based tools. Nucleic Acids Res 2013;41:D590–6.

Sekiguchi Y, Yamada T, Hanada S et al. Anaerolinea thermophila gen. nov., sp. nov. and Caldilinea aerophila gen. nov., sp. nov., novel filamentous thermophiles that represent a previously uncultured lineage of the domain Bacteria at the subphylum level. Int J Syst Evol Microbiol 2003;53:1843–51.

Seviour EM, Blackall LL, Christensson C et al. The filamentous morphotype Eikelboom Type 1863 is not asingle genetic entity. J Appl Microbiol 1997;82:411–21.

Sorokin DY, Lücker S, Vejmelkova D et al. Nitrification expanded: discovery, physiology and genomics of a nitrite-oxidizing bacterium from the phylum Chloroflexi. ISME J 2012;6:2245–56.

Speirs L, Nittami T, McIlroy S et al. Filamentous bacterium Eikelboom type 0092 in activated sludge plants in Australia is a member of the phylum Chloroflexi. Appl Env Microbiol 2009;75:2446–52.

Speirs LBM, Dyson ZA, Tucci J et al. Eikelboom filamentous morphotypes 0675 and 0041 embrace members of the Chloroflexi: resolving their phylogeny, and design of fluorescence in situ hybridisation probes for their identification. FEMS Microbiol Ecol 2017;93, DOI: 10.1093/femsec/fix115.

Speirs LBM, McIlroy SJ, Petrovski S et al. The activated sludge bulking filament Eikelboom morphotype 0914 is a member of the Chloroflexi. Env Microbiol Rep 2011;3:159–65.

Speirs LBM, Tucci J, Seviour RJ. The activated sludge bulking filament Eikelboom morphotype 0803 embraces more than one member of the *Chloroflexi*. Stams A (ed.). FEMS Microbiol Ecol 2015;91:fiv100.

Streichan M, Golecki JR, Schon G. Polyphosphate-accumulating bacteria from sewage plants with different proceses for biological phosphorus removal. FEMS Microbiol Lett 1990;73:113–24.

Wallner G, Amann R, Beisker W. Optimizing fluorescent in situ hybridization with rRNA-targeted oligonucleotide probes for flow cytometric identification of microorganisms. Cytometry 1993;14:136–43.

Wanner J, Kragelund C, Nielsen PH. Microbiology of bulking. In: Seviour RJ, Nielsen PH (eds.). Microbial Ecology of Activated Sludge. London: IWA Publishing, 2010, 191–214.

Xia Y, Kong Y, Nielsen PH. In situ detection of protein-hydrolysing microorganisms in activated sludge. FEMS Microbiol Ecol 2007;60:156–65.

Yamada T, Imachi H, Ohashi A et al. Bellilinea caldifistulae gen. nov., sp. nov. and Longilinea arvoryzae gen. nov., sp. nov., strictly anaerobic, filamentous bacteria of the phylum Chloroflexi isolated from methanogenic propionate-degrading consortia. Int J Syst Evol Microbiol 2007;57:2299–306.

Yamada T, Sekiguchi Y, Hanada S et al. Anaerolinea thermolimosa sp. nov., Levilinea saccharolytica gen. nov., sp. nov. and Leptolinea tardivitalis gen. nov., sp. nov., novel filamentous anaerobes, and description of the new classes Anaerolineae classis nov. and Caldilineae classis nov. in the. Int J Syst Evol Microbiol 2006;56:1331–40.

Yilmaz LS, Parnerkar S, Noguera DR. mathFISH, a web tool that uses thermodynamics-based mathematical models for in silico evaluation of oligonucleotide probes for fluorescence in situ hybridization. Appl Env Microbiol 2011;77:1118–22.

Ziegler AS, McIlroy SJ, Larsen P et al. Dynamics of the Fouling Layer Microbial Community in a Membrane Bioreactor. PloS One 2016;11:e0158811.

